# Genetically diverse populations spread faster in benign but not in challenging environments

**DOI:** 10.1101/2020.11.27.400820

**Authors:** Mortier Frederik, Masier Stefano, Bonte Dries

## Abstract

Population spread from a limited pool of founding propagules is at the basis of biological invasions. The size and genetic variation of these propagules eventually affect whether the invasion is successful or not. The inevitable bottleneck at introduction decreases genetic diversity, and therefore should affect population growth and spread. However, many heavily bottlenecked invasive populations have been successful in nature. These negative effects of a genetic bottleneck are typically considered to be relaxed in benign environments because of a release from stress. Despite its relevance to understand and predict invasions, empirical evidence on the role of genetic diversity in relation to habitat quality is largely lacking. We use the mite *Tetranychus urticae* Koch as a model to experimentally assess spread rate and the size of genetically depleted inbred populations and enriched mixed populations. This was assessed in replicated linear patch systems consisting of benign (bean), challenging (tomato) or a gradient (bean to tomato) habitat. We find that genetic diversity increased population spread rates in the benign but not in the challenging habitat. Additionally, variance in spread was consistently higher in genetically poor populations and highest in the challenging habitat. Our experiment challenges the general view that a bottleneck in genetic variation decreases invasion success in challenging, but not in benign environments.

## Introduction

Environmental change can move the physiological limits of a range, and therefore lead to range expansions as determined by population growth and spread (Chuang and Peterson 2016). Ranges can alternatively expand beyond the existing geographical limits by the introduction of individuals away from their original range. But alongside the environmental opportunities for range expansions, population spread requires the individual capabilities to do so. Individual-level life history traits related to reproduction and dispersal will influence the extent and variation in population spread and therefore range border dynamics (Fisher 1937, Angert et al. 2011). As these traits have a genetic basis in many organisms (Roff 2001, Saastamoinen et al. 2018), range dynamics should to an important extent be determined by the population’s genetic composition. Genetic diversity, in numbers and in variation in identity of genotypes, has a well-studied positive effect on various ecological processes. Genetic diversity tends to improve ecological performance as expressed by fitness associated proxies as higher population growth rates, productivity (e.g. Reusch et al. 2005) or movement (e.g. Wagner et al. 2017). This positive relationship between genetic diversity and a variety of demographic processes can be explained by several mechanisms (Hughes et al. 2008, Bolnick et al. 2011).

1. A higher genetic diversity increases opportunities for natural selection to act, hence increasing the average fitness in the population which can eventually increase population growth and improve overall ecological performance. Other evolutionary processes, like inbreeding depression, may in contrast decrease ecological performance.
2. Enhanced sampling in genetically diverse populations increases the probability of the presence of a phenotype with a positive impact on ecological performance.
3. A higher genetic diversity increases the variance in phenotypes which can result in an increase of the mean population’s ecological an improved ecological performance relative to the average phenotype when a convex relationships exist between genetic diversity and the ecological function (Jensen’s inequality principle).
4. Complementarity effects like niche partitioning and facilitation increase ecological performance by diversifying the ways of performing well.
5. Portfolio effect stabilize fluctuations in the ecological function. Fluctuations from different genotypes that differ in frequency combine to less fluctuating dynamics.

Range dynamics are even more strongly affected by genetic diversity at longer evolutionary time scales: phenotypes may organize themselves along the range resulting in more dispersive phenotypes disproportionally closer to the leading range edge (Phillips et al. 2010, Burton et al. 2010, Phillips and Perkins 2019). This spatial sorting and subsequent spatial selection is known to accelerate range expansion (Fronhofer and Altermatt 2015, Szűcs et al. 2017, Van Petegem et al. 2018). Genetic drift during spread may, however, slow expansion (Peischl et al. 2015). These evolutionary processes additionally influence the variability in range expansion in both deterministic and stochastic ways (Williams et al. 2019).

Spread during an biological invasion fundamentally differs from spread from an established range. At introduction, the invading population’s genetic diversity is constrained by several bottlenecks during its transport from the ancestral population to form the eventual founding population (Pierce et al. 2017, Renault et al. 2018). The number of founders and the potential admixture of different founding lines (Dlugosch and Parker 2008, Rius and Darling 2014) eventually determines the severity of this genetic bottleneck. Some invasive populations even show a higher diversity than their ancestral counterparts (Estoup et al. 2016). The number of invaders also imposes demographic effects. A smaller invading population is more vulnerable to Allee effects (Stephens et al. 1999, Taylor and Hastings 2005) and demographic stochasticity (Fauvergue et al. 2012), and is therefore less likely to be successful. The number of invaders is often found to increase invasion success (Colautti et al. 2006, Simberloff 2009, Blackburn et al. 2015). And while the different impact of genetic diversity and demography at establishment has been studied (Ahlroth et al. 2003, Szűcs et al. 2014, Vahsen et al. 2018, Sinclair et al. 2019), its importance for the subsequent population spread is not yet resolved (but see Wagner et al. 2017). Such insights are especially needed to solve the genetic paradox of invasions, i.e. the success of invasions despite severe reductions of genetic diversity (Dlugosch and Parker 2008, Mullarkey et al. 2013, Estoup et al. 2016, Schrieber and Lachmuth 2017).

The environment in which the population spreads is another important ecological driver of invasion success. Introductions can occur in an environment that is similar or vastly different from their ancestral one. When the environment of introduction is different and challenging, evolutionary rescue by means of adaptation (Bell and Gonzalez 2011) offers a possible route to tackle the imposed challenges and stimulate population growth and spread. In a benign environment, such adaptations may not be needed to attain high population sizes (Schrieber and Lachmuth 2017). Inbreeding depression is also stronger manifested in a challenging and stressful environment (Fox and Reed 2011). Decreased genetic diversity may therefore constrain population spread in stressful but not in benign environments. The difference in importance of genetic diversity for establishment between benign and challenging environments has already been experimentally demonstrated (Hufbauer et al. 2013, Szűcs et al. 2014) and is hinted at in some natural invasions (Daehler and Strong 1997, Hawley et al. 2005). Environments are, however, seldom homogenous in the environmental parameters that determine a species’ niche. Rather they gradually change in an autocorrelated way (Legendre 1993). Such environmental gradients are anticipated to affect the rate and success of spread in ways that are different from homogenous benign and challenging environments. Like for evolutionary processes (Bell and Gonzalez 2011), a gradual increase in stress may favor population spread compared to a sudden change into a challenging environment and enforce an ecological rescue mechanism.

The impact of evolution on range expansion dynamics in different environments is unpredictable (Williams et al. 2019), but significant insights in range expansion can be gained by studying the immediate effects of genetic diversity on the onset of expansions. The genetic diversity of founders can inform us which populations are initially better primed to expand their range, to get ahead and ultimately to invade successfully. We specifically expect genetic diversity to increase population spread more in a challenging environment compared to a benign one. We also explored the effect of genetic diversity on variability in population spread. In parallel with Williams et al. (2019), the direction of this effect likely depends on the balance of deterministic and stochastic forces, affecting the predictability of its effect. Because failed invasions cannot be studied in nature, we established replicated experimental populations of *Tetranychus urticae* Koch (two-spotted spider mite) in linear patch systems in which population spread rate and demography were followed for approximately three generations. Spreading populations varied in their level of genetic diversity at the start (single-female lines or mixed lines), independent of the number of introduced individuals, and differed in the kind of environment they were introduced to (benign, challenging or a gradient from benign to challenging).

## Methods

### Model system

We tested population spread of *Tetranychus urticae* Koch (two-spotted spider mite), a generalist arthropod herbivore. This mite is a known pest species that has been found on more than 900 hosts all over the world (Navajas 1998). The species is a model in ecological and evolutionary research because of its ease of use, high fertility and annotated genome (Belliure et al. 2010, Macke et al. 2011, Van Petegem et al. 2018, Masier and Bonte 2020). For this experiment, we collected twelve natural populations from a variety of host plants and two lab-populations in September 2018 (more information in Appendix S1). We sampled at least 50 individuals with many more in most of the sampled populations. We maintained the collected populations on *Phaseolus vulgaris* (bean) leaf patches on wet cotton in petri dishes (150 mm diameter) sealed off by a lid with ventilation through which the mites could not escape to keep populations from contaminating each other. We maintained these populations in the lab for six months, which amounted to around fifteen generations, before the start of the first procedures (creating the mixed lines, see below).

We used *Phaseolus vulgaris* var. prelude (bean) plants or leaf patches to keep our stocks on, to perform the single female line procedures and as a benign host in the experiment. Bean is known to be an optimal host that shows little defense against the mites. We used *Solanum lycopersicum* moneymaker (tomato) as a challenging host. The mites occur on tomato but in past experiments, they attained a lower fitness on it (0.2-0.25 of fecundity on bean, Alzate et al. 2017). Local adaptation to tomato is possible but is never observed to result in a higher fitness on tomato compared to bean (Alzate et al. 2017, Mortier and Bonte 2020). None of the natural populations were sampled from tomato or a host that is taxonomically from the same family (Solanaceae).

### Genetic diversity

To obtain starting populations of mites with different levels of genetic variation, we created one genetically rich population by mixing all collected wild and laboratory lines. Carbonnelle et al. (2007) demonstrated significant genetic differences between natural populations of T. urticae in Western Europe at similar geographic scales. The genetically rich mix was formed two months prior to the start of the experiment, spanning around five generations, in order to leave the mixed population enough time to avoid any effect of outbreeding depression. We kept this mixed line in four crates with four to eight bean plants each that were regularly mixed. Each bean plant contained a few hundred to a thousand individuals at all times resulting in a total population of the order of magnitude of ten thousand. This setup supported a high population size to avoid subsequent loss of genetic diversity due to drift or effects of linkage disequilibrium as much as possible. Multiple genetically poor populations were established as single female lines from eight of the collected wild and laboratory lines. The single-female lines were formed by sampling one unfertilized female from a collected line by transferring one quiescent deutonymph, the mite’s life stage on the verge of adulthood, to a separate leaf. Due to their parthenogenetic nature, unfertilized eggs from that mite will exclusively produce males that are, then, back-crossed with their mother. As a result, the female produces fertilized eggs hatching males and females. With this procedure, we established a population that consists only of genetic material from the original female, not considering mutations. Low genetic diversity is better retained by smaller population sizes on the leave disks compared to the whole plants. Since densities have been shown to provoke maternal effects on dispersal distance (Bitume et al. 2014), we ensured the populations on whole plants and on leaf discs to be maintained at the same high densities, close to carrying capacity. We performed this procedure with four unfertilized female from each collected line to hedge for likely failure. A mother laid eggs and the unfertilized eggs developed at 26°C, close to the maximum developmental speed. The mother was kept at 17°C while males developed in order to slow the mother’s ageing and preserve her fertility to the time of fertilization. During this procedure and in the following four weeks to the start of the experiment, we kept the mites at all times on bean patches (±5cm×6cm) on wet cotton in a petri dish.

### Population spread

We tested population spread dynamics in experimental linear system containing plant patches that were connected with bridges (Appendix S2: fig. S2). Every experimental population was placed in a clean plastic crate (26.5cm × 36.5cm) covered in three layers of wet cotton wool (Rolta®soft). Patches of plant leaves (1.5cm × 2.5cm) were connected with one another by a parafilm® bridge (1cm × 8cm) touching the leaf patch, with the remaining edge aligned with paper towel strips. The wet cotton provided an impenetrable and deadly matrix in between plant patches and provided the cost of moving from one patch to another in the form of mortality risk. We started each population spread test by sampling 40 individuals from a start population and placing them on the first patch in their population spread arena. We added two additional connected patches. Every day we added, if needed, new connected patches to always have two empty patches in front of the furthest occupied patch and every two days we replaced the still unoccupied patches to keep the unoccupied patches at the front fresh and attractive for potential spreading mites. From past experience with similar patch setups, we know that mites very rarely disperse more than two patches per day under the established densities. Additionally, all plant patches were replaced with a fresh patch once a week. The old patch was placed upside down on the fresh patch for two days in order for mites of all life stages to move from to the fresh patch. Because the old patch always dried quickly, most mites moved to the fresh patch within those two days. Replacing all plant patches replenished resources to sustain the core of the population. These linear patch systems snaked through our crates for twelve possible patches (Appendix S2: fig. S2). In case a thirteenth (and subsequent) patch was needed, the first (and subsequent) patch and its connection to the next was removed to make space for the new one. We sacrificed trailing patches since we were mostly interested in the leading edge dynamics. The arenas were kept at room temperature, around 23°C, with a 16:8h L:D photoperiod.

We started population spread tests in three environments: 1) a benign environment of all bean patches, 2) a challenging environment of all tomato patches and 3) a gradient from benign to challenging patches (with 3 bean patches, 1 tomato patch, 2 bean patches, 2 tomato patches, 1 bean patch, 3 tomato patches, 1 bean patch followed by all tomato patches). We started sixteen population spread tests in each environment. Eight tests of a genetically poor population were each started from a single female line from a different natural population. Eight tests of a genetically rich population were each started from mites from a different plant in the mixed population. We recorded population spread as the furthest occupied patch in each range on a daily basis and we recorded the number of mites on each patch on a weekly basis over the duration of 35 days or five weeks. Though generations started to overlap, we estimate that this amounted to around three generations.

### Statistical models

We analyzed the outcome of our experiment using Bayesian inference. In R (3.6.3), the ‘brms’ (2.12.0; Bűrkner 2018) package implements ‘Stan’ (Carpenter et al. 2017) as a framework for parameter posterior estimation using Hamiltonian Monte Carlo (HMC).

#### Mean population spread

We constructed a multi-level model to estimate effects on mean population spread. We modelled furthest occupied patch as response variable with a Gaussian distribution. We modelled a fixed effect of time, the environmental treatment and the genetic diversity treatment and a variable intercept and slope (in time) for each population spread arena (i.e. random effect of replicate population spread arena and its interaction with time).

#### Variability in population spread

We estimated effects on variance in population spread among replicates from the same treatment for each point in time. We again modelled population spread (furthest occupied patch) with a Gaussian distribution, but estimated effects on both the mean and standard deviation. We modelled the mean of the Gaussian distribution with a fixed effect of time, the environmental treatment and the diversity treatment. However, we modelled no variable intercepts but pooled that variation around the mean originating from among different replicates together with the residual variation in order to model all variation around the mean. We then proceeded to model the standard deviation of the Gaussian distribution also with a fixed effect of time, the environmental treatment in and the diversity treatment in the same model. This way, the model estimated the standard variation for each treatment at each point in time. We calculated coefficients of variance from the estimated mean and standard deviation in population spread.

#### Total population size

Population size can help explain population spread. Therefore, we modelled total population size across all patches as a response variable with a negative binomial distribution. We model a fixed effect of time, the environmental treatment and the diversity treatment and a variable intercept and slope (in time) for each population spread arena.

#### Population density

We modelled total population sizes across all patches as a response variable with a negative binomial distribution. We model a fixed effect of the amount of occupied patches, the environmental treatment and the diversity treatment. We build a model like the previous but with the amount of occupied patches as fixed effect to understand whether larger populations were larger because they occupied more patches or because they had a higher density.

We report model outcomes as plots of the posterior distribution or direct calculations of marginal effects of the posterior distribution. Bayesian inference considers less the most likely parameter or marginal effect value but rather the whole distribution of likely values, which can most faithfully be reported visually instead of as a metric. We always plotted the 0.09 and 0.91 percentiles of the likelihood distribution. We want to stress that this is by no means a significance threshold, but only serves as two arbitrary extremes to aide interpretation of the plotted distribution. We also include the estimated difference between two groups differing in a treatment to directly assess the estimated effect of that treatment in most plots.

A more detailed description of the statistical models and their outcomes can be found in the supplementary materials (see Appendix S3). The data files and scripts to analyze them are published (doi: 10.5281/zenodo.4025183).

## Results

### Mean population spread

All populations showed an initial burst of spread during the first days where the relative high density of mites at the starting patch incentivized mites to leave for the next patch. This resulted in an estimated intercept higher than one, the expected intercept as all populations started with one occupied patch (fig. 1, top). Mites in the tomato environment spread very little after this initial burst resulting in the edge lingering around the fourth patch on average. In the bean and gradient environment, however, the population on average kept spreading for the duration of the experiment. The higher genetic diversity of the mixed populations resulted in a faster spreading population in the bean environment as seen in the predominantly negative estimated difference in slope between both diversity treatments (sfl-mix, fig. 1, bottom left). Genetic diversity had no convincing effect in the gradient or tomato environment as seen in the estimated differences in slope (sfl-mix) around zero (fig. 1, bottom middle, bottom right). On bean, mixed lines reached on average the ninth patch while single female lines reached on average the sixth patch. On the gradient, both treatments reached on average the eighth patch. Contrary to expectations, the effect of genetic diversity on population spread was larger in the benign relative to the challenging environment.

**Figure 1:**
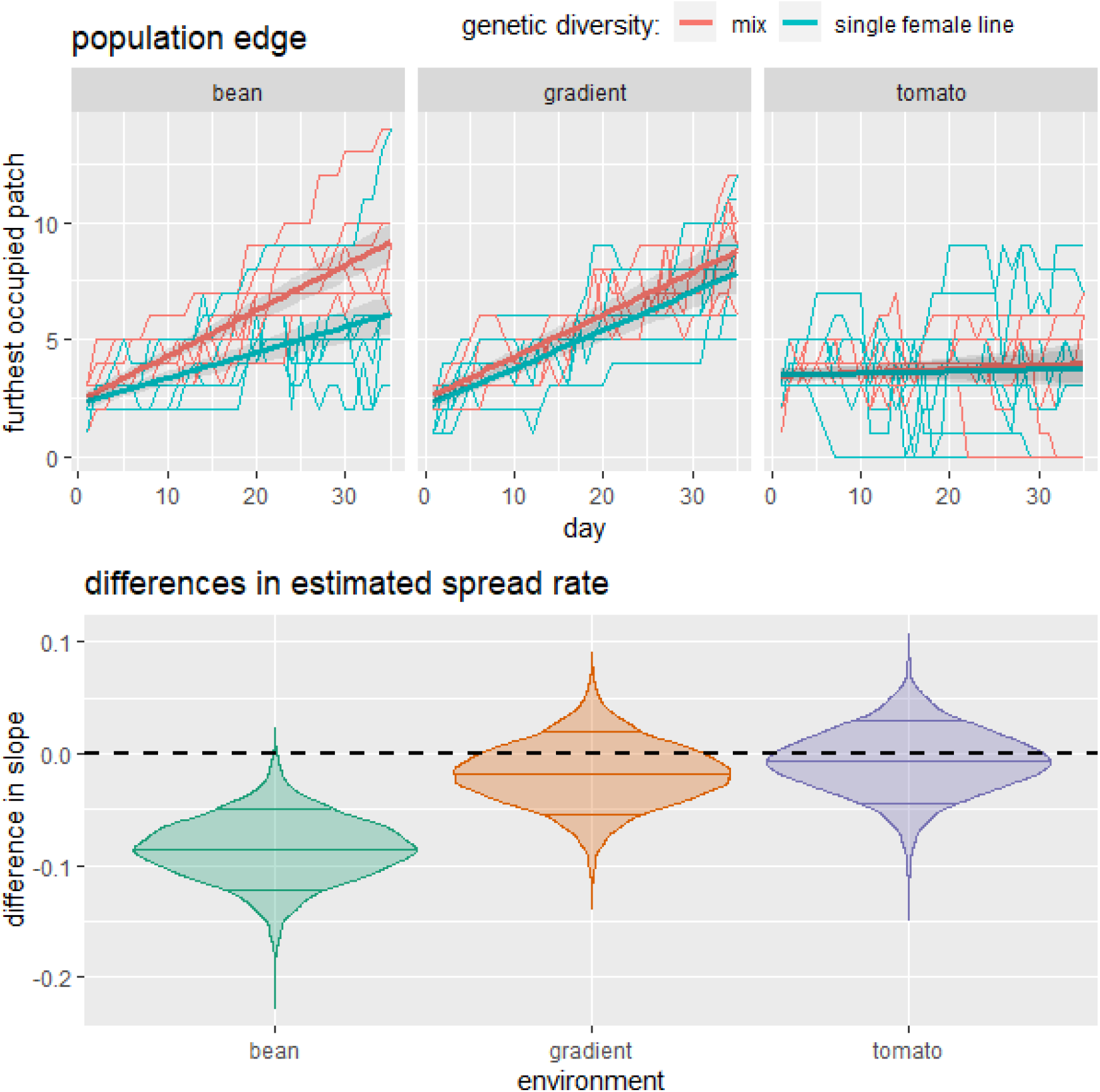
Top: population spread as the furthest occupied patch over the duration of the experiment for mixed (red) and single female (blue) populations in the bean (left), gradient (middle) or tomato (right) environment. The fine lines show the recorded spread of each population for each day while the wide lines with shades represent the statistical (BMC) model estimate with likelihood interval of the 0.09 and 0.91 quantiles. Bottom: differences in estimated slopes of population spread in time (single female line - mix) in the bean (left), gradient (middle) or tomato (right) environment. The dashed line indicates equal estimated slopes.

### Variability in population spread

The genetically rich mixed lines had a less variable population spread compared to the single female lines in all environments (fig. 2). In the bean and gradient environments, variance was more or less constant and the difference in variance due to genetic diversity was relatively small. However in the tomato environment, the coefficient of variance increased in time and show a relatively sizeable difference in the coefficient of variance between genetically poor and rich lines.

**Figure 2:**
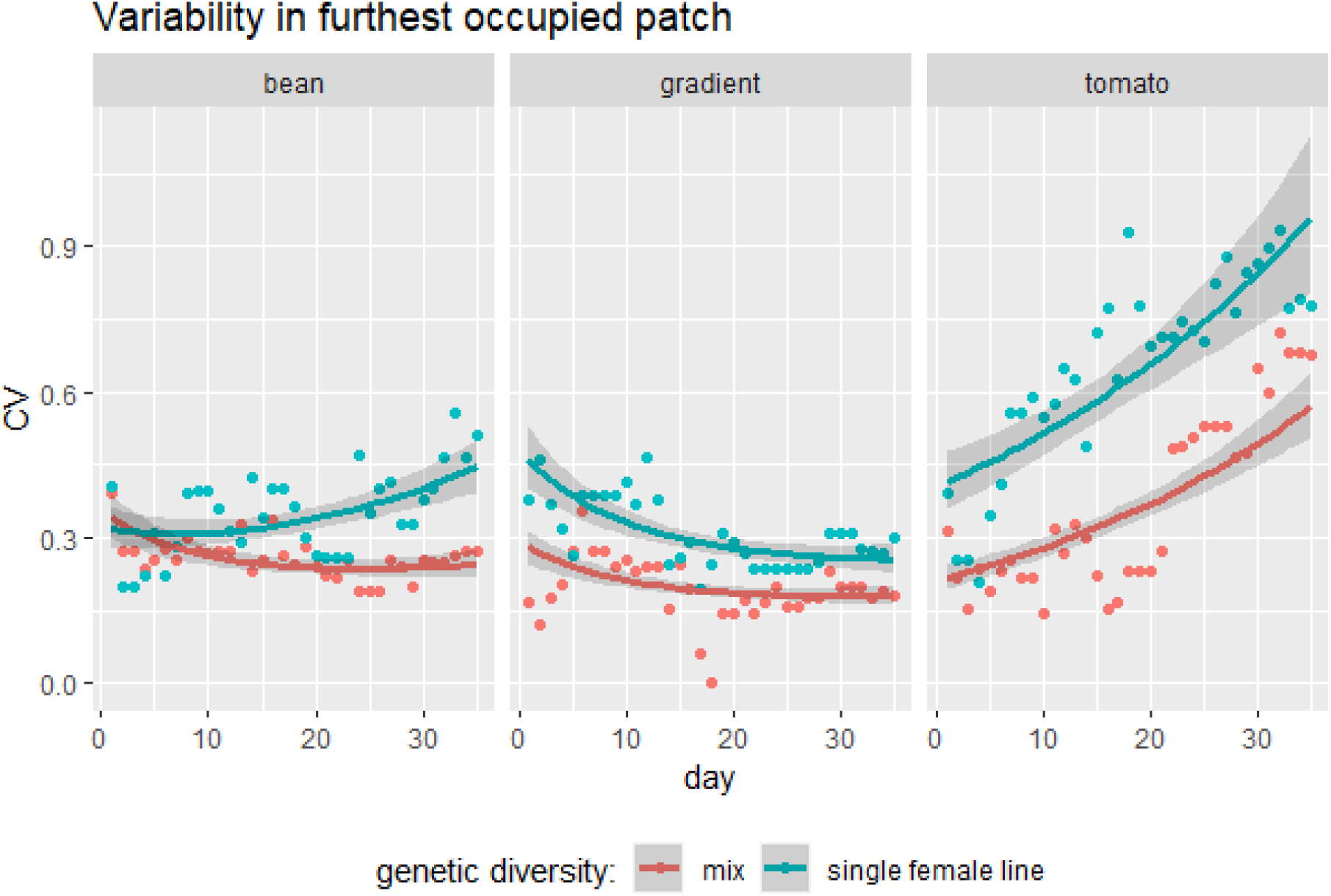
population spread variability as coefficient of variance (CV) of the number of occupied patches over the duration of the experiment for mixed (red) and single female (blue) populations in the bean (left), gradient (middle) or tomato (right) environment. The dots show the calculated coefficient of variance in population spread (standard deviation in population spread divided by mean population spread) among all populations each day while the wide lines with shades represent the statistical (BMC) model estimate with likelihood interval of the 0.09 and 0.91 quantiles

### Total population size

Total population sizes were smaller on tomato compared to the bean and gradient treatment (fig. 3, top). On tomato we found a larger population size of mixed compared to single female lines (fig. 3, bottom right). The estimated differences between the genetically diverse and depleted lines on bean had zero (i.e. the no difference), close to the 91 percentile visual aide we plot (fig. 3, left). We estimated no difference on the gradient (fig. 3, bottom middle).

**Figure 3:**
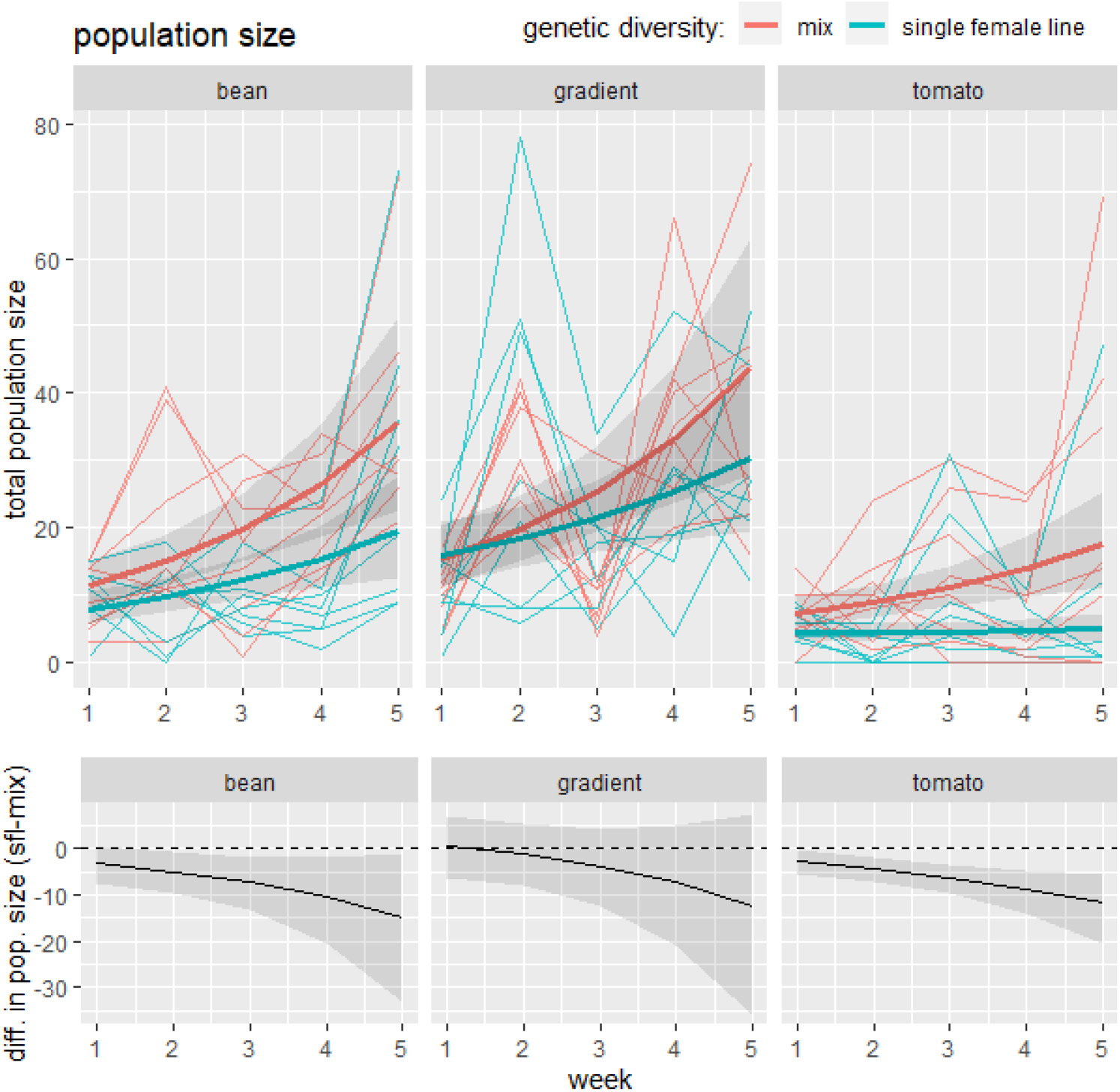
total population size (top) for mixed (red) and single female (blue) populations and estimated differences in total population size between single female lines and mixes (sfl-mix) (bottom) in the bean (left), gradient (middle) or tomato (right) environment. The fine lines show the recorded total population size of each population each week while the wide lines with shades represent the statistical (BMC) model estimate with likelihood interval of the 0.09 and 0.91 quantiles. The estimated differences in total population size (bottom) between single female lines and mixes (sfl-mix) visualizes whether the differences due to genetic diversity in its respective top panel differs from zero (dashed line).

### Population density

By adding the number of occupied patches as a predictor to the model instead of time, we found total population size to be affected by the amount of patches: total population size increased with plant patches (fig.4, top). Population sizes for a given number of plant patches of mixed and single female lines were estimated not to differ on bean and the gradient (fig. 4, bottom left, middle). However, mixed lines showed a higher population size for a given number of plant patches on tomato, as shown by the increasingly positive difference in population size (fig. 4, bottom right). This indicated a higher density in mixed lines than in single female lines. Densities of many mixed lines on tomato even exceeded densities in the benign environment.

**Figure 4:**
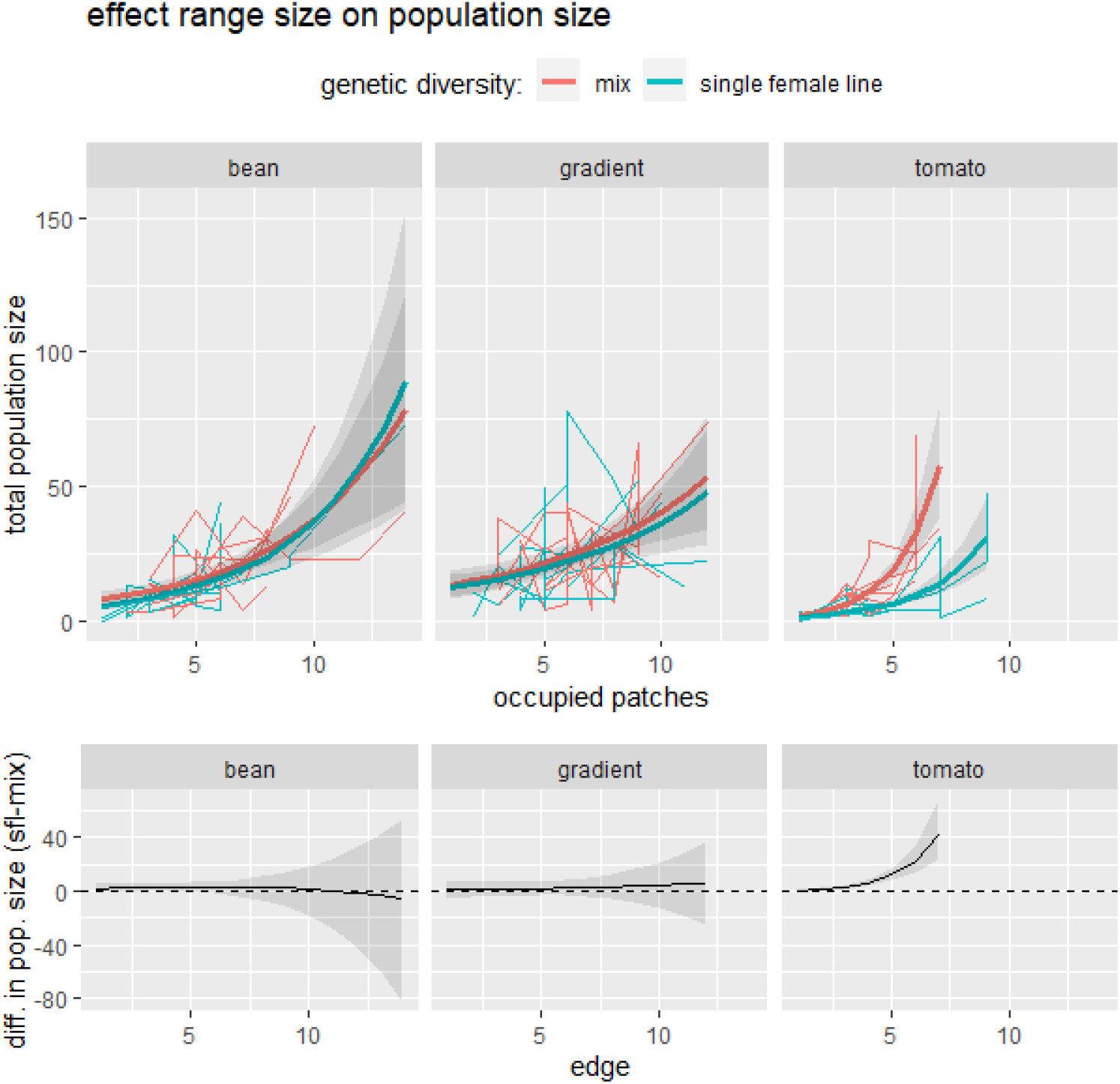
total population size regressed over the furthest occupied patch by that population (top) for mixed (red) and single female (blue) populations and estimated differences in total population size between single female lines and mixes (sfl-mix) (bottom) in the bean (left), gradient (middle) or tomato (right) environment. The fine lines show the recorded total population size of each population each week while the lines and shades represent the statistical (BMC) model estimate with confidence likelihood of the 0.09 and 0.91 quantiles. The estimated differences in total population size (bottom) between single female lines and mixes (sfl-mix) visualizes whether the differences due to genetic diversity in its respective top panel differs from zero (dashed line). All recorded time points are plotted for every population and the model only estimates the range of occupied patches that was observed per combination of diversity and environment.

## Discussion

The hypothesis of genetic diversity benefiting population size and spread in challenging but not in benign environments is only partially validated by our experiment. In contrast to common predictions, we document a positive effect of genetic diversity on population spread rate in the benign bean environment but not in the challenging tomato environment. In support, we detected a positive effect of genetic diversity on total and local population sizes in the challenging tomato environment, but not in the benign bean environment. This accords with findings of Hufbauer et al. (2013) and Szucs et al. (2014) for population growth in whiteflies and flour beetles.

In line with Wagner et al. (2017), we found that diversity accelerated spread in the benign environment. Such increased expansions should theoretically result from increased population growth and dispersal (Fisher 1937, Angert et al. 2011), or any combination of these two. Diversity effects are known to affect population growth (Hughes et al. 2008), and can lead to a fast population growth to or even beyond carrying capacity. At this population size. A positive density-dependence of dispersal (Matthysen 2005, Bowler and Benton 2005, Kawasaki et al. 2017) then, can circumvent strong density regulation by competition as a result from strong growth and lead to a faster range expansions (‘so-called pushed range expansions; Dahirel et al. 2020). Average local population sizes in our experiment did not differ conclusively between the diversity treatments, whereas those of spread did. We therefore attribute the accelerated range expansion in part to the fact that diverse population may impose higher per capita dispersal rates by for instance a sampling effect that enables more dispersive genotypes to be present or heterosis and positive body condition effects on dispersal (Wagner et al. 2017).

In contrast to the benign environment, genetically diverse and less diverse populations both stop expanding after the initial leap in spread in the challenging environment. This means that less diverse populations spread as fast as diverse populations under these conditions, while reaching lower population sizes. Negative effects of a low genetic diversity on reproduction are therefore anticipated to be compensated by an increase in dispersal. Dispersal to avoid kin competition is well documented in *Tetranychus urticae* (Bitume et al. 2013, Van Petegem et al. 2018) and is a plausible cause of the enhanced dispersal in genetically impoverished populations. Mixed and single female lines reach equal high densities in the benign environment and any kin-competition is likely overruled by resource competition which reverses a negative effect of diversity on dispersal to a positive.

Population dynamics on the gradient share characteristics from both the homogeneous benign bean and homogeneous challenging tomato environments. Ranges on the gradient reached on average the position where the mites started to encounter more tomato patches than bean patches, with some populations spreading beyond this point. But because population spread started in the benign environment, populations on the gradient unsurprisingly grew and spread at similar rates to populations on bean. No diversity effects were however observed. The genetically poor populations spread as far as the enriched populations on the gradient, but further than genetically poor populations on bean. We therefore attribute the spread of genetically poor populations to the environment rather than to a diversity effect. The insertion of challenging patches likely stimulated dispersing mites to skip the challenging patches in search of the next bean patch, which suggest an additional effect of the gradient’s patchy nature. Obviously, informed movement leading to habitat choice must enable such a behavior (Egas and Sabelis 2001, Mortier and Bonte 2020).

The impact of spatial sorting, selection and local adaptation on range expansions, have been studied experimentally in recent years (Fronhofer and Altermatt 2015, Van Petegem et al. 2016, 2018, Szűcs et al. 2017). We deliberately limited our experiment to roughly 2-3 generations to focus on the immediate effect of genetic diversity on the initial population spread. Selection as a precursor of local adaptation and spatial evolution may play an important role from the first generation onwards (Szűcs et al. 2017). All replicate populations spread at least a few patches and assuming some heritable difference in the ability to spread, this should result in some spatial sorting while determining the resulting population spread rate at the same time. Similarly, the ability to better reproduce in the local environment, if heritable, will be proportionally overrepresented in the next generation and may increase population spread rate simultaneously. It is therefore clear that if such mechanisms, or any other long term mechanism, would have manifested during the short term of the experiment, we are actually underestimating them.

Genetically diverse lines were characterized by a lower variability in spread among replicates in every environment, hence showing a more consistent and predictable spread rate. This consequence of genetic diversity is found for many ecological processes (Hughes et al. 2008). The difference between genetically diverse and poor populations is larger and starts earlier in time on the challenging tomato. Williams et al. (2019) attribute variability in range expansion dynamics to the balance of variance generating (stochastic) and variance reducing (deterministic) evolutionary forces. Analogously, we propose that different diversity effects can have a variance increasing or decreasing effect on population spread. A probably relevant effect is that higher population sizes reduce demographic stochasticity as a consequence. Since movement is positively related to density, and hence population size, variability in spread is also expected to be minimized in genetically diverse populations. This explains differences in variability between the diversity treatments on tomato and explains the higher overall variability on tomato compared to the other environments. We furthermore speculate that larger differences in variability between genetically diverse and poor replicates on tomato are due to specific genotypes disproportionally impacting range spread. Population spread in the genetically poor lines will be either high or low depending on the presence or absence of the impactful genotype. In comparison, all genetically rich lines are more likely to contain this or any other high impact genotype, hence reducing any variance in population spread among replicates. The smaller difference in variability between genetically diverse and poor lines in the benign bean environment suggests a smaller contribution of specific high-dispersive genotypes, but rather a larger contribution of positive effects of diversity in and of itself. Our data does not allow us to infer a specific mechanism but we speculate that niche partitioning of residents and dispersers is at play. We are not talking about niche partitioning in terms of type of resource but in terms of diverging movement strategies to exploit the same limited resources in the landscape (Bonte et al. 2014, Schlägel et al. 2020). It is more expected that competitive subordinates move as an adaptive behavior when competition is strong, as expected in a benign environment (stabilizing mechanisms; cfr. Chesson 2000). Such fitness stabilizing mechanisms can accelerate range spread in genetically diverse populations, not because of the presence of a specific genotype but because the diversity of genotypes provoking synergies that cannot be reached in isolation. It is not unlikely that a possible higher diversity of movement strategies may have induced some strategies to disperse more to outrun competition in the benign environment.

The view that invaded environments are the ones where species are released from stress is one of the most important paradigms in invasion biology. The enemy-release hypothesis, for instance, builds on this view of invaded environments being inherently benign (Keane 2002, Colautti et al. 2004). Such a mechanism implies that the release of stressful interactions should outweigh any potential maladaptation to the new environment. We do not see any reason why this assumption should hold. Rather, a strong bias exists in invasion biology as all observed invasions have been by definition successful, with any reference to failed invasions in both challenging and benign environments lacking. Experiments like ours that observe aspects of ongoing invasion, spread in our case, instead of the aftermath, circumvents this survivorship bias. Another aspect of the classic view on invasions is that high propagule pressure, which avoids demographic stochasticity and results in a higher genetic diversity, increases an invader’s success. Our results from challenging environments however illustrates that a genetic bottleneck does not decrease population spread in the initial phase and, therefore, may ultimately not decrease invasion success. This is opposite to the established explanation for the genetic paradox of invasions that a genetic bottleneck should still affect a population in a challenging environment. However, our results still support the idea that the effect of a genetic bottleneck is conditional on the environment. Bear in mind that these challenging environments already support a lower invasion success inherently that, as we showed, may not be aided by genetic diversity. Genetic adaptation to the challenging environment may, however impose a niche shift on the longer term, and therefore allowing the invading population to capitalize on its genetic diversity. Such niche shifts have been observed in experimental (Szűcs et al. 2017) and successful natural invasions (Broennimann et al. 2007), but tend to be the exception rather than the rule (Peterson 2011, Petitpierre et al. 2012, Strubbe et al. 2013).

## Acknowledgements

FM thanks the Special Research Fund (BOF) of Ghent University for a PhD scholarship. SM thanks Fonds Wetenschappelijk Onderzoek (FWO) of Flanders for a PhD scholarship. DB, FM and SM are additionally supported by FWO research grant G018017N. We also thank Nicky Wybouw from the department of plants and crops (UGent) for providing the two laboratory population of tomato-bred two-spotted spider mites. Lastly, we thank Steven Goossens with help in collecting some of the natural mite populations.

