## Appendix S1 for "Genetically diverse populations spread faster in benign but not in challenging environments"

Journal: Ecology

### Appendix 1: sample locations:

Information on the mite populations used in the study. We provide coordinates from where the natural populations were sampled. This is not relevant for the two lab populations. We also note the host plant the population was sampled from. Populations indicated in blue were used to form their single-female line. All populations were mixed to become the mixed population as described in the main text.

| name | Coordinates/location | Host plant |
| --- | --- | --- |
| 1 | N51.0038° E3.8083° | *Chelidonium majus* (greater celandine) |
| 2 | N51.0348° E3.7232° | unknown |
| 3 | N51.0350° E3.7228° | *Aristolochia fimbriata* |
| Greenhouse (GH) | N51.0354° E3.7228° | *Cucumis sativus* (cucumber) |
| Oost-Duinkerke (OD) | N51.1246° E2.6833° | *Humulus lupulus* (common hop) |
| De Haan (DH) | N51.2823° E3.0538° | *Euonymus europaeus* (spindle) |
| Knokke | N51.3509° E3.3360° | *Sambucus nigra* (black elder) |
| Steven | N51.0333° E4.1525° | *Phaseolus vulgaris* (bean) |
| Melle | N51.0046° E3.8024 | *Lonicera nitida* (box honeysuckle) |
| Citadel | N51.0377° E3.7231° | *Lonicera nitida* (box honeysuckle) |
| Flora | N51.0300° E3.7436° | *Chelidonium majus* (greater celandine) |
| Bourgoyen (Bour) | N51.0661° E3.6806 | *Chelidonium majus* (greater celandine) |
| SR-VL | lab | *Phaseolus vulgaris* (bean) |
| MR-VL | lab | *Phaseolus vulgaris* (bean) |
