## Appendix S2 for "Genetically diverse populations spread faster in benign but not in challenging environments"

Journal: Ecology

### Appendix S2: range spread arena diagram

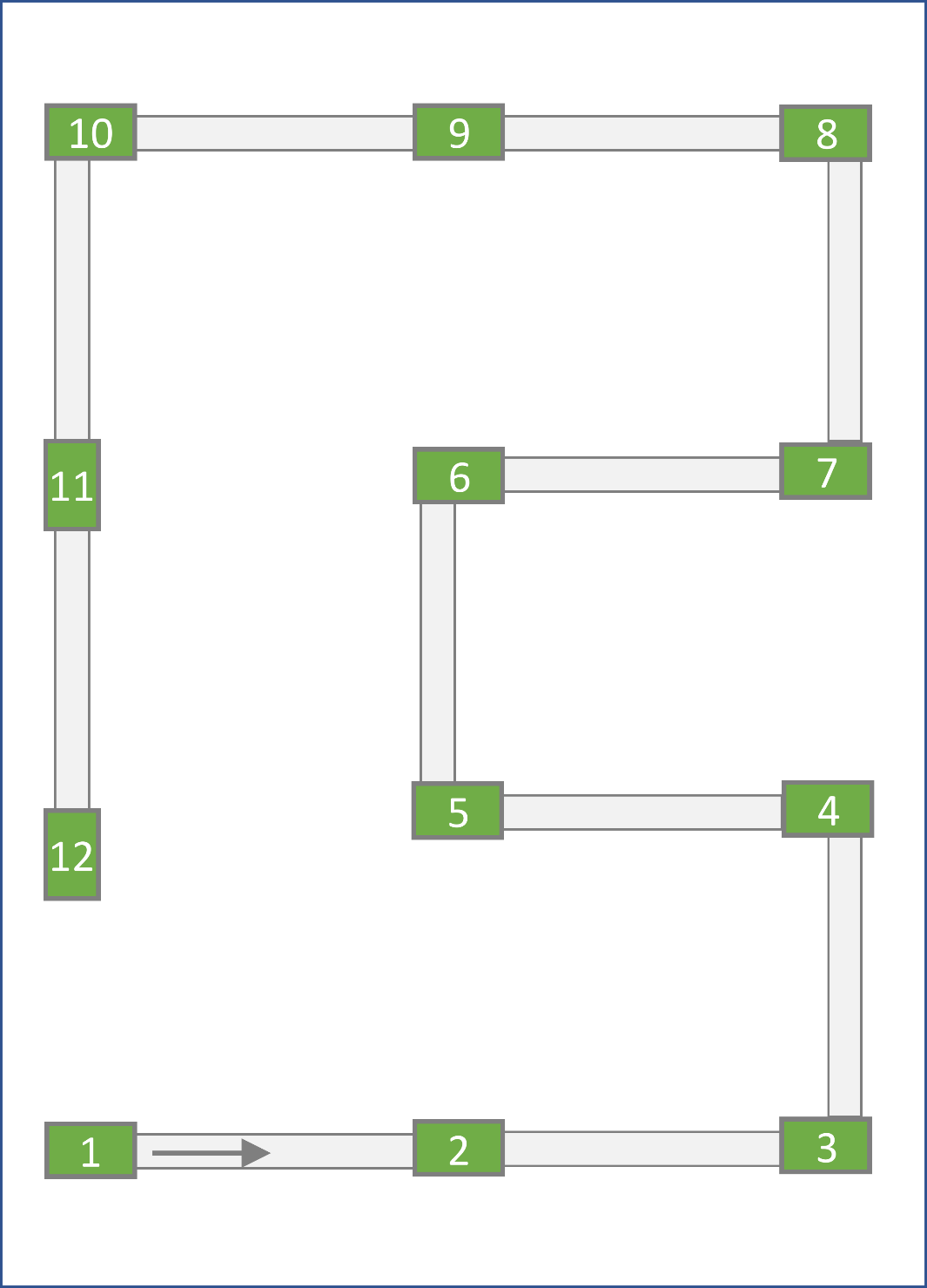

### *Figure S1: range spread arena that consists of plant patches (green rectangles) that are connected to each other by parafilm® bridges (grey rectangles) in a linear sequence. Mites are introduced at the starting patch (upper right).*
