## Appendix S3 for "Genetically diverse populations spread faster in benign but not in challenging environments"

Journal: Ecology

### Appendix 3: Statistical model description and estimates

All models were run in R (3.6.3) using brms (2.12.0) and Hamiltonian Monte Carlo with two chains with each 5000 iteration from which 2000 were warmup.

Data files and scripts to analyze them are published (doi: 10.5281/zenodo.4025183).

Model 1: Mean population spread

a) Model representation

edge ~ normal(1 + environment*day*diversity + (1 + day|line))

prior = c(prior(normal(0,4), class = Intercept),

prior(normal(0,4), class = b),

prior(cauchy(0,2), class = sd),

prior(cauchy(0, 2), class = σ),

prior(lkj(2), class = cor))

With *edge* the furthest occupied patch of a breeding *line* of a certain *diversity* spreading in a certain *environment* recorded at a certain point in time (in number of *day*s since the start). We modelled edge with a normally distributed error distribution and estimated it depending on *environment*, time (*day*) and *diversity* and all their interactions. We also modelled a varying intercept and slope in time for each tested breeding *line*. We used priors that are weakly regularizing. Of note, the used LKJ distribution is a Lewandowski-Kurowicka-Joe distribution that is regularly used a prior for a correlation matrix.

b) Model parameter estimates

```
                      mean se_mean   sd     2.5%     97.5% n_eff Rhat
b_Int                 2.34    0.00 0.22     1.91     2.78  3111    1
b_envG                0.10    0.00 0.25    -0.39     0.58  3119    1
b_envT                1.09    0.00 0.24     0.61     1.57  3244    1
b_day                 0.19    0.00 0.02     0.16     0.24  1937    1
b_divsfl             -0.09    0.01 0.31    -0.71     0.52  2897    1
b_envG:day           -0.01    0.00 0.01    -0.04     0.01  3328    1
b_envT:day           -0.18    0.00 0.01    -0.20    -0.16  3246    1
b_envG:divsfl        -0.22    0.01 0.35    -0.90     0.46  3044    1
b_envT:divsfl         0.13    0.01 0.34    -0.53     0.80  3156    1
b_day:divsfl         -0.08    0.00 0.03    -0.14    -0.03  2340    1
b_envG:day:divsfl     0.07    0.00 0.02     0.03     0.10  2891    1
b_envT:day:divsfl     0.08    0.00 0.02     0.05     0.11  3090    1
sd_line__Int          0.37    0.00 0.13     0.13     0.66  2389    1
sd_line__day          0.05    0.00 0.01     0.03     0.08  1904    1
cor_line__Int__day   -0.25    0.01 0.28    -0.73     0.35  1299    1
σ                     1.44    0.00 0.03     1.39     1.49 11781    1
r_line[2,Int]        -0.01    0.00 0.25    -0.52     0.49  7031    1
r_line[Bourg,Int]    -0.21    0.00 0.25    -0.75     0.28  6819    1
r_line[Citadel,Int]   0.27    0.00 0.27    -0.23     0.82  4977    1
r_line[Flora,Int]     0.03    0.00 0.25    -0.45     0.53  6579    1
r_line[GH,Int]        0.04    0.00 0.25    -0.45     0.54  5390    1
r_line[OD,Int]       -0.27    0.00 0.26    -0.82     0.22  6069    1
r_line[S1,Int]        0.58    0.01 0.30     0.03     1.21  3195    1
r_line[S2,Int]        0.16    0.00 0.26    -0.34     0.69  5291    1
r_line[S3,Int]       -0.07    0.00 0.25    -0.57     0.43  5925    1
r_line[S4,Int]        0.11    0.00 0.25    -0.37     0.60  6657    1
r_line[S5,Int]       -0.24    0.00 0.26    -0.78     0.23  5785    1
r_line[S6,Int]       -0.02    0.00 0.24    -0.52     0.46  7013    1
r_line[S7,Int]        0.02    0.00 0.25    -0.48     0.51  6138    1
r_line[S8,Int]       -0.53    0.00 0.29    -1.14    -0.02  4190    1
r_line[SRVL,Int]      0.15    0.00 0.25    -0.32     0.66  5797    1
r_line[Steven,Int]   -0.01    0.00 0.25    -0.51     0.49  6949    1
r_line[2,day]        -0.05    0.00 0.02    -0.09    -0.01  3198    1
r_line[Bourg,day]     0.00    0.00 0.02    -0.04     0.04  3304    1
r_line[Citadel,day]   0.07    0.00 0.02     0.03     0.11  3347    1
r_line[Flora,day]    -0.04    0.00 0.02    -0.08     0.01  3119    1
r_line[GH,day]       -0.07    0.00 0.02    -0.11    -0.03  2936    1
r_line[OD,day]        0.00    0.00 0.02    -0.04     0.04  3103    1
r_line[S1,day]       -0.06    0.00 0.02    -0.11    -0.02  2305    1
r_line[S2,day]       -0.05    0.00 0.02    -0.09    -0.01  2213    1
r_line[S3,day]        0.02    0.00 0.02    -0.03     0.06  2515    1
r_line[S4,day]        0.01    0.00 0.02    -0.04     0.05  2535    1
r_line[S5,day]        0.02    0.00 0.02    -0.02     0.07  2519    1
r_line[S6,day]        0.01    0.00 0.02    -0.03     0.05  2489    1
r_line[S7,day]        0.03    0.00 0.02    -0.01     0.07  2498    1
r_line[S8,day]        0.03    0.00 0.02    -0.02     0.07  2551    1
r_line[SRVL,day]      0.04    0.00 0.02     0.00     0.08  3170    1
r_line[Steven,day]    0.05    0.00 0.02     0.01     0.09  3184    1
```

c) Model outcome

As presented in the main text, we observed no difference in mean population spread between mixed and single-female lines in the gradient and tomato environment, but we found a convincing higher spread for mixed lines in the bean environment (fig. 1, top). This convincing difference clearly stemmed from a difference in slopes (fig 1, middle) and not from a difference in intercepts (fig. 1, bottom). While all population spread tests started at the first patch, we did not fix the modelled intercept at 1 to enable the model to capture a possible early burst in spread.

Model 2: Population spread variance

a) Model representation

edge ~ normal(μ ~ 1 + environment*day*diversity, σ ~ 1 + day*environment*div)

prior = c(prior(normal(0,4), class = Intercept),

prior(normal(0,4), class = b)

Here, we model the population spread as a distributional model that estimates the effects of predictors on the mean, but also the standard deviation of spread. We model population spread with a normal error distribution, a mean population spread depending on *environment*, time (*day*) and *diversity* and their interactions; and standard deviation depending on *environment*, time (*day*) and *diversity* and their interactions. We use weakly regularizing priors. The priors that we give above are used for intercepts and coefficients of predictors for the linear sub model in both mean and standard deviation.

b) Model parameter estimates

```
                      mean se_mean   sd     2.5%     97.5% n_eff Rhat
b_Int                 2.45    0.00 0.13     2.18     2.70  2430    1
b_σ_Int              -0.14    0.00 0.09    -0.31     0.04  1720    1
b_envG                0.05    0.00 0.17    -0.28     0.38  2467    1
b_envT                0.85    0.00 0.18     0.50     1.20  2674    1
b_day                 0.19    0.00 0.01     0.17     0.21  2511    1
b_divsfl             -0.32    0.00 0.18    -0.67     0.02  2166    1
b_envG:day           -0.01    0.00 0.01    -0.03     0.01  2440    1
b_envT:day           -0.16    0.00 0.01    -0.19    -0.14  2897    1
b_envG:divsfl        -0.06    0.01 0.26    -0.56     0.44  2414    1
b_envT:divsfl         0.55    0.01 0.30    -0.06     1.15  2827    1
b_day:divsfl         -0.07    0.00 0.01    -0.09    -0.04  2416    1
b_envG:day:divsfl     0.06    0.00 0.02     0.03     0.09  2489    1
b_envT:day:divsfl     0.05    0.00 0.02     0.01     0.09  2962    1
b_σ_day               0.03    0.00 0.00     0.02     0.03  1637    1
b_σ_envG             -0.18    0.00 0.12    -0.41     0.06  1830    1
b_σ_envT             -0.22    0.00 0.12    -0.46     0.02  1822    1
b_σ_divsfl           -0.24    0.00 0.13    -0.48     0.01  1573    1
b_σ_day:envG          0.00    0.00 0.01    -0.02     0.01  1778    1
b_σ_day:envT          0.01    0.00 0.01     0.00     0.02  1758    1
b_σ_day:divsfl        0.01    0.00 0.01     0.00     0.03  1561    1
b_σ_envG:divsfl       0.58    0.00 0.17     0.25     0.92  1636    1
b_σ_envT:divsfl       0.95    0.00 0.18     0.60     1.29  1708    1
b_σ_day:envG:divsfl  -0.02    0.00 0.01    -0.03     0.00  1567    1
b_σ_day:envT:divsfl  -0.02    0.00 0.01    -0.04    -0.01  1756    1
```

c) Model outcome

We calculated fitted values for mean and standard deviation for each posterior sample and calculate the estimated coefficient of variance of population spread by dividing estimated standard deviation by estimated mean. By calculating coefficients of variance, there is no intercept or slope estimated.

We found a lower coefficient of variance in population spread of mixed lines compared to single-female lines consistently over all environments (fig. S2). We also noticed a clearly bigger difference in the tomato environment, on which the variance increases clearly with time.

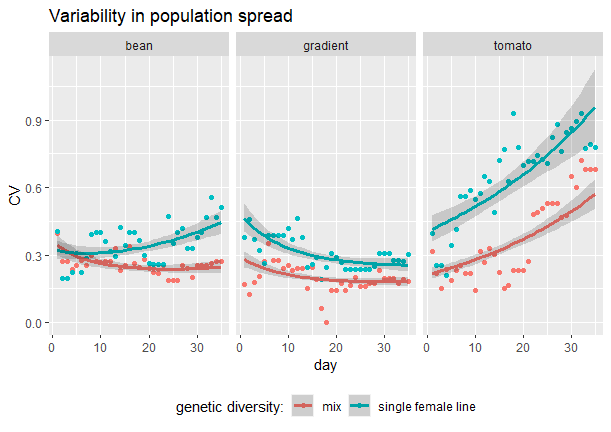

Figure S2: population spread variability as coefficient of variance (CV) of the number of sequential patches the population occupies over the duration of the experiment for mixed (red) and single female (blue) populations in the bean (left), gradient (middle) or tomato (right) environment. The plotted points show the recorded spread of each population each day while the lines and shades represent the statistical (BMC) model estimate with confidence interval of the 0.09 and 0.91 quantiles

Model 3: Total population size

a) Model representation

total population size ~ negbinom(1 + environment*week*diversity + (1+week|line))

prior = c(prior(normal(0,4), class = Intercept),

prior(normal(0,4), class = b),

prior(cauchy(0,2), class = sd),

prior(lkj(2), class = cor),

prior(cauchy(0,2), class = shape)),

With *total population size* the counted amount of mites (larvae, nymphs and adults) of a population with a certain genetic *diversity* (mixed line or single-female line) in a certain *environment* (bean, gradient or tomato) at a point in time (*week*). We modelled a negative binomial error distribution of the counted total population size and estimated an intercept, the linear effect of these three predictors and all their interactions. We also modelled a varying intercept and slope in time (*week*) for each tested breeding *line*. We used priors that are weakly regularizing.

b) Model estimates

```
                      mean se_mean   sd    2.5%   97.5% n_eff Rhat
b_Int                 2.12    0.01 0.29    1.54   2.69  2814    1
b_envG                0.35    0.01 0.40   -0.42   1.14  2765    1
b_envT               -0.35    0.01 0.41   -1.15   0.46  2763    1
b_week                0.28    0.00 0.09    0.10   0.46  2451    1
b_divsfl             -0.28    0.01 0.40   -1.04   0.52  2730    1
b_envG:week          -0.03    0.00 0.12   -0.27   0.20  2925    1
b_envT:week          -0.07    0.00 0.12   -0.31   0.17  2634    1
b_envG:divsfl         0.38    0.01 0.56   -0.73   1.47  2689    1
b_envT:divsfl        -0.06    0.01 0.58   -1.24   1.06  2622    1
b_week:divsfl        -0.06    0.00 0.13   -0.32   0.18  2586    1
b_envG:week:divsfl   -0.03    0.00 0.17   -0.35   0.31  2714    1
b_envT:week:divsfl   -0.12    0.00 0.18   -0.45   0.24  2534    1
sd_line__Int          0.17    0.00 0.13    0.01   0.47  2318    1
sd_line__week         0.11    0.00 0.04    0.03   0.20  1206    1
cor_line__Int__week   0.00    0.01 0.42   -0.76   0.80   973    1
shape                 1.99    0.00 0.24    1.56   2.50  5479    1
r_line[2,Int]        -0.07    0.00 0.19   -0.52   0.28  2187    1
r_line[Bourg,Int]    -0.09    0.00 0.18   -0.58   0.20  4334    1
r_line[Citadel,Int]   0.12    0.01 0.23   -0.25   0.70  1469    1
r_line[Flora,Int]    -0.04    0.00 0.16   -0.42   0.28  4590    1
r_line[GH,Int]       -0.03    0.00 0.18   -0.43   0.34  2647    1
r_line[OD,Int]        0.04    0.00 0.16   -0.28   0.42  5786    1
r_line[S1,Int]       -0.04    0.00 0.17   -0.43   0.29  2680    1
r_line[S2,Int]       -0.10    0.00 0.19   -0.59   0.20  2734    1
r_line[S3,Int]       -0.04    0.00 0.17   -0.44   0.28  5394    1
r_line[S4,Int]       -0.02    0.00 0.17   -0.42   0.31  5784    1
r_line[S5,Int]        0.02    0.00 0.16   -0.31   0.37  4508    1
r_line[S6,Int]        0.10    0.00 0.19   -0.20   0.58  2580    1
r_line[S7,Int]        0.03    0.00 0.16   -0.28   0.40  5352    1
r_line[S8,Int]        0.05    0.00 0.16   -0.23   0.45  4182    1
r_line[SRVL,Int]      0.05    0.00 0.17   -0.26   0.44  3002    1
r_line[Steven,Int]    0.05    0.00 0.16   -0.25   0.45  5442    1
r_line[2,week]       -0.12    0.00 0.08   -0.30   0.02  2750    1
r_line[Bourg,week]   -0.02    0.00 0.07   -0.16   0.13  5028    1
r_line[Citadel,week]  0.18    0.00 0.09    0.01   0.37  2050    1
r_line[Flora,week]   -0.03    0.00 0.07   -0.17   0.10  4760    1
r_line[GH,week]      -0.11    0.00 0.08   -0.28   0.04  3177    1
r_line[OD,week]       0.00    0.00 0.07   -0.15   0.14  5335    1
r_line[S1,week]      -0.09    0.00 0.08   -0.25   0.05  3168    1
r_line[S2,week]      -0.08    0.00 0.08   -0.24   0.07  3556    1
r_line[S3,week]       0.03    0.00 0.07   -0.10   0.18  4678    1
r_line[S4,week]       0.04    0.00 0.07   -0.09   0.19  4899    1
r_line[S5,week]       0.03    0.00 0.07   -0.11   0.17  4910    1
r_line[S6,week]       0.09    0.00 0.08   -0.06   0.24  3298    1
r_line[S7,week]      -0.01    0.00 0.07   -0.15   0.13  5376    1
r_line[S8,week]       0.01    0.00 0.07   -0.14   0.15  5142    1
r_line[SRVL,week]     0.08    0.00 0.07   -0.05   0.23  3893    1
r_line[Steven,week]   0.02    0.00 0.07   -0.11   0.17  5624    1
```

c) Model outcome

As presented in the main text, we found an unconvincing difference in mean population size and population increase between genetically diverse and less diverse lines in the bean and gradient environment but we found a convincingly higher mean population size of mixed lines in the tomato environment (fig. S3, top). When plotting the estimated differences in intercepts and the estimated differences in slopes between mixed and single-female lines in each environment, we noticed a slight but not clear difference in slopes and intercepts for the tomato environment (fig. S3, middle, bottom). A relatively strong negative correlation, then, explained how unconvincing differences in slopes and estimates resulted in a convincing overall difference in population sizes. Posterior samples where mixed lines did not have a higher intercept than single female lines on tomato were very likely to have a higher slope and samples where mixed lines did not have a higher slope than single female lines on tomato were very likely to have a higher intercept (fig. S4).

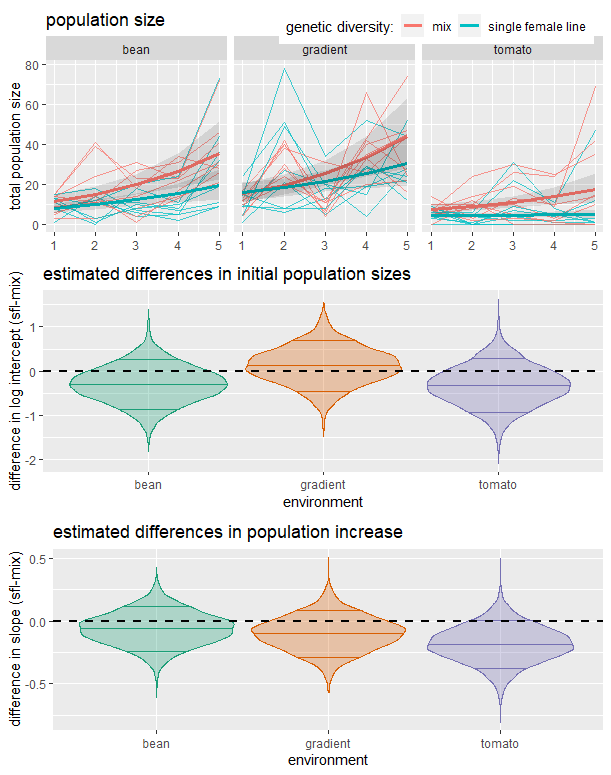

Figure S3: Top: total population size for mixed (red) and single female (blue) populations in the bean (left), gradient (middle) or tomato (right) environment. The fine lines show the recorded total population size of each population each week while the lines and shades represent the statistical (BMC) model estimate with likelihood interval of the 0.09 and 0.91 quantiles. Middle: differences in estimated intercepts of total population size (single female line - mix) in the bean (left), gradient (middle) or tomato (right) environment. Bottom: differences in estimated slopes of total population size in time (single female line - mix) in the bean (left), gradient (middle) or tomato (right) environment. The dashed line indicates equal estimated intercepts and slopes respectively.

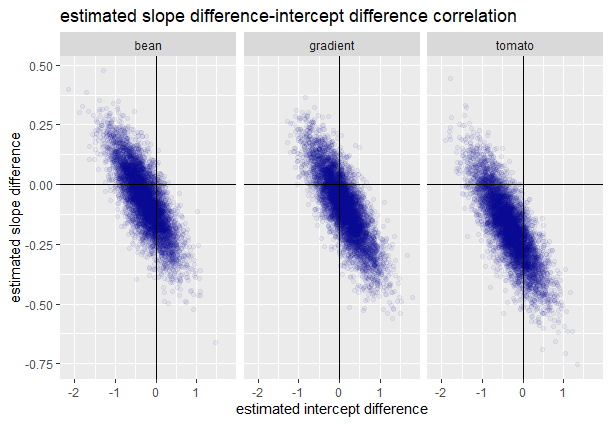

Figure S4: correlation of estimated differences in slope and differences in intercept of total population size regressed over time (single female line - mix) in the bean (left), gradient (middle) or tomato (right) environment. The vertical and horizontal black line indicates equal estimated intercepts and equal estimated slopes respectively.

Model 4: Range size effect on total population size

a) Model representation

Total population size ~ negbinom(1 + environment*edge*diversity)

prior = c(prior(normal(0,4), class = Intercept),

prior(normal(0,4), class = b),

prior(cauchy(0,2), class = sd),

prior(cauchy(0,2), class = shape))

With *total population size* the counted amount of mites (larvae, nymphs and adults) of a population with a certain genetic *diversity* (mixed line or single-female line) in a certain *environment* (bean, gradient or tomato) at a point in time (*week,* not modelled here). We modelled a negative binomial error distribution of the counted total population size and estimated an intercept, the linear effect of *diversity*, *environment*, the amount of occupied patches (*edge*) and all their interactions. We also modelled a varying intercept for each tested breeding *line*. We used priors that are weakly regularizing.

b) Model estimates

```
                     mean se_mean   sd    2.5%    97.5% n_eff Rhat
b_Int                1.84    0.01 0.29    1.28    2.43  2088    1
b_envG               0.57    0.01 0.43   -0.26    1.44  2107    1
b_envT              -1.70    0.01 0.43   -2.53   -0.87  1876    1
b_edge               0.18    0.00 0.04    0.09    0.26  2155    1
b_divsfl            -0.33    0.01 0.38   -1.10    0.42  1802    1
b_envG:edge         -0.05    0.00 0.06   -0.18    0.08  2068    1
b_envT:edge          0.38    0.00 0.09    0.21    0.56  2210    1
b_envG:divsfl        0.25    0.01 0.57   -0.88    1.37  2234    1
b_envT:divsfl        0.07    0.01 0.55   -1.01    1.14  1791    1
b_edge:divsfl        0.03    0.00 0.06   -0.09    0.15  1855    1
b_envG:edge:divsfl  -0.03    0.00 0.09   -0.22    0.15  2276    1
b_envT:edge:divsfl  -0.19    0.00 0.12   -0.42    0.03  1989    1
sd_line__Int         0.19    0.00 0.08    0.04    0.37  1643    1
shape                2.88    0.01 0.36    2.24    3.66  4874    1
r_line[2,Int]       -0.14    0.00 0.15   -0.48    0.13  4150    1
r_line[Bourg,Int]   -0.07    0.00 0.14   -0.35    0.20  5595    1
r_line[Citadel,Int]  0.16    0.00 0.15   -0.09    0.50  3773    1
r_line[Flora,Int]    0.01    0.00 0.14   -0.26    0.30  6261    1
r_line[GH,Int]      -0.12    0.00 0.15   -0.44    0.15  5240    1
r_line[OD,Int]       0.13    0.00 0.14   -0.12    0.44  3796    1
r_line[S1,Int]      -0.08    0.00 0.14   -0.38    0.18  4912    1
r_line[S2,Int]      -0.12    0.00 0.15   -0.43    0.14  4898    1
r_line[S3,Int]      -0.04    0.00 0.14   -0.34    0.22  5218    1
r_line[S4,Int]      -0.11    0.00 0.14   -0.42    0.15  3915    1
r_line[S5,Int]       0.06    0.00 0.13   -0.20    0.34  5328    1
r_line[S6,Int]       0.22    0.00 0.16   -0.03    0.55  2828    1
r_line[S7,Int]      -0.03    0.00 0.14   -0.32    0.23  5254    1
r_line[S8,Int]       0.11    0.00 0.14   -0.14    0.43  3798    1
r_line[SRVL,Int]     0.11    0.00 0.15   -0.15    0.43  4694    1
r_line[Steven,Int]  -0.09    0.00 0.14   -0.40    0.17  5332    1
```

c) Model outcome

As presented in the main text, we found no difference in mean population size for a certain amount of occupied patches between genetically diverse and less diverse lines in the bean and gradient environment but we found convincingly bigger populations for a certain amount of occupied patches in the tomato environment (fig. S3, top). Plotting the estimated differences in intercepts and the estimated differences in slopes between mixed and single-female lines in each environment, we noticed almost no difference in intercepts and a noticeable-but-not-definitive difference in slopes for the tomato environment (fig. S5, middle, bottom). A relatively strong negative correlation, then, explained why the overall difference in population sizes was more convincing than the difference in slopes. The few posterior samples where mixed lines did not have a higher slope than single female lines on tomato were very likely to have a higher intercept and samples where mixed lines did not have a higher intercept than single female lines on tomato were very likely to have a higher slope (fig. S6).

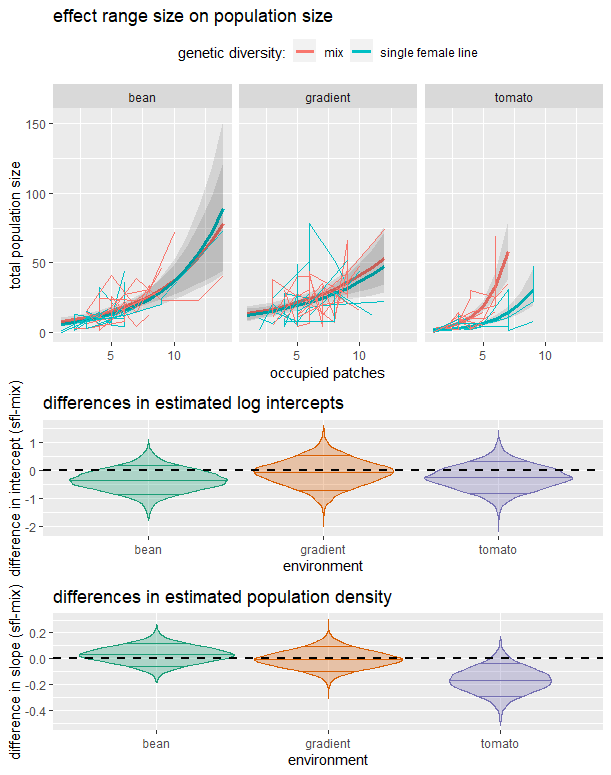

Figure S5: Top: total population size regressed over the furthest occupied patch by that population for mixed (red) and single female (blue) populations in the bean (left), gradient (middle) or tomato (right) environment. The fine lines show the recorded total population size of each population each week while the lines and shades represent the statistical (BMC) model estimate with likelihood interval of the 0.09 and 0.91 quantiles. Middle: differences in estimated intercepts of total population size (single female line - mix) in the bean (left), gradient (middle) or tomato (right) environment. Bottom: differences in estimated slopes of total population size regressed over furthest occupied patch (single female line - mix) in the bean (left), gradient (middle) or tomato (right) environment. The dashed line indicates equal estimated intercepts and slopes respectively.

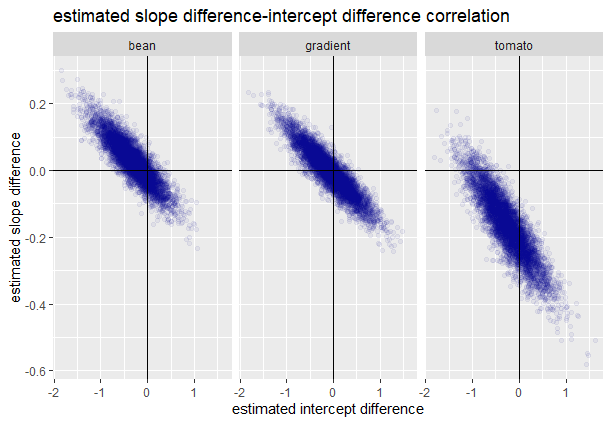

Figure S6: correlation of estimated differences in slope and differences in intercept of total population size regressed over furthest occupied patch (single female line - mix) in the bean (left), gradient (middle) or tomato (right) environment. The vertical and horizontal black line indicates equal estimated intercepts and equal estimated slopes respectively.
